# Rapid and Sensitive Protein Complex Alignment with Foldseek-Multimer

**DOI:** 10.1101/2024.04.14.589414

**Authors:** Woosub Kim, Milot Mirdita, Eli Levy Karin, Cameron L.M. Gilchrist, Hugo Schweke, Johannes Söding, Emmanuel D. Levy, Martin Steinegger

## Abstract

Advances in computational structure prediction will vastly augment the hundreds of thousands of currently-available protein complex structures. Translating these into discoveries requires aligning them, which is computationally prohibitive. Foldseek-Multimer computes complex alignments from compatible chain-to-chain alignments, identified by efficiently clustering their superposition vectors. Foldseek-Multimer is 3-4 orders of magnitudes faster than the gold standard, while producing comparable alignments; allowing it to compare billions of complex-pairs in 11 hours. Foldseek-Multimer is open-source software: github.com/steineggerlab/foldseek, webserver: search.foldseek.com and the BFMD database.

---

The similarity between two protein complexes is reflected in their optimal structural alignment, which also dictates a pairing of their chains. Aligning and comparing quaternary structures is essential for quantifying their structural diversity and identifying structural similarities and changes across different conformations or homologs. Furthermore, it is important to understanding protein function because many proteins operate as complexes (1).

Recently, Foldseek (2) has been developed as a fast structural aligner to detect similarity between two single-chain proteins, expressed using 3Di, a designated alphabet for describing tertiary amino acid interactions. Using Foldseek allows searching for similar single-chain structures in large databases, such as the AFDB (3). However, since aligning two complexes requires knowing the correct pairing of their chains, Foldseek cannot be used directly to find the alignment between them. US-align (4) is a structural aligner for various types of molecules, including protein complexes. Its strategy for complex alignment is TM-score maximization. As there is a factorial number of possible assignments of chain pairings, US-align employs a greedy search heuristic for proposing candidate assignments, which are refined by dynamic programming. This heuristic was shown to make US-align up to five times faster than the state-of-the-art MM-align (5), while producing higher scoring alignments, making US-align the gold-standard for pairwise complex alignment.

Aiming to discover pairs of structurally conserved interfaces in large databases, Dey et al. developed QSalign (6) for the detection of similar homomeric complexes. QSalign saves computation time by performing the full pairwise structural alignment only on complex pairs prefiltered based on their sequence similarity, retaining pairs with ca. 25% sequence identity or more. This speed-up comes at the expense of sensitivity, limiting its ability to discover structurally similar pairs in the twilight zone or below. Despite this speed-up, QSalign still took several months to conduct an all-vs.-all search encompassing about a hundred thousand complexes in the 3DComplex DB V5 (7) using 100 threads. An alternative approach to reduce computational time during database search was presented by Guzenko et al. (8), who compare the shapes between two complexes through 3D Zernike descriptors, avoiding the need to pair their chains. This approach can query through hundreds of thousands of structures in less than a second. However, it can only discover global matches between molecules of similar shapes, limiting its sensitivity, compared to chain-pairing methods, like US-align and QSalign. Furthermore, it is unable to find local matches within chains that do not match globally.

The challenge of sensitively searching large databases is expected to intensify as the computational prediction of protein complexes using tools like AlphaFold-Multimer (9) can now be performed on entire proteomes to systematically predict complexes (10–12) and on sequences from metagenomic samples. This will enrich our databases with a plethora of structures, potentially in the millions, in the coming years.

To address the need for large-scale structural comparisons between complexes, we developed Foldseek-Multimer (**Fig. 1**). Three factors contribute to its speed: 1) using Foldseek for fast chain-to-chain comparison, 2) describing chain-to-chain alignments as superposition vectors, and using them to identify complex alignments by efficient clustering, and 3) utilizing clustered databases during search. Through benchmarks, we show that Foldseek-Multimer is: 1) nearly as accurate as US-align, while being orders of magnitude faster, 2) sensitive and suitable for metagenomic studies of complexes with low sequence similarity to others, 3) capable of all-vs.-all searches, examining billions of complex-pairs in 11h.

**Fig. 1.**
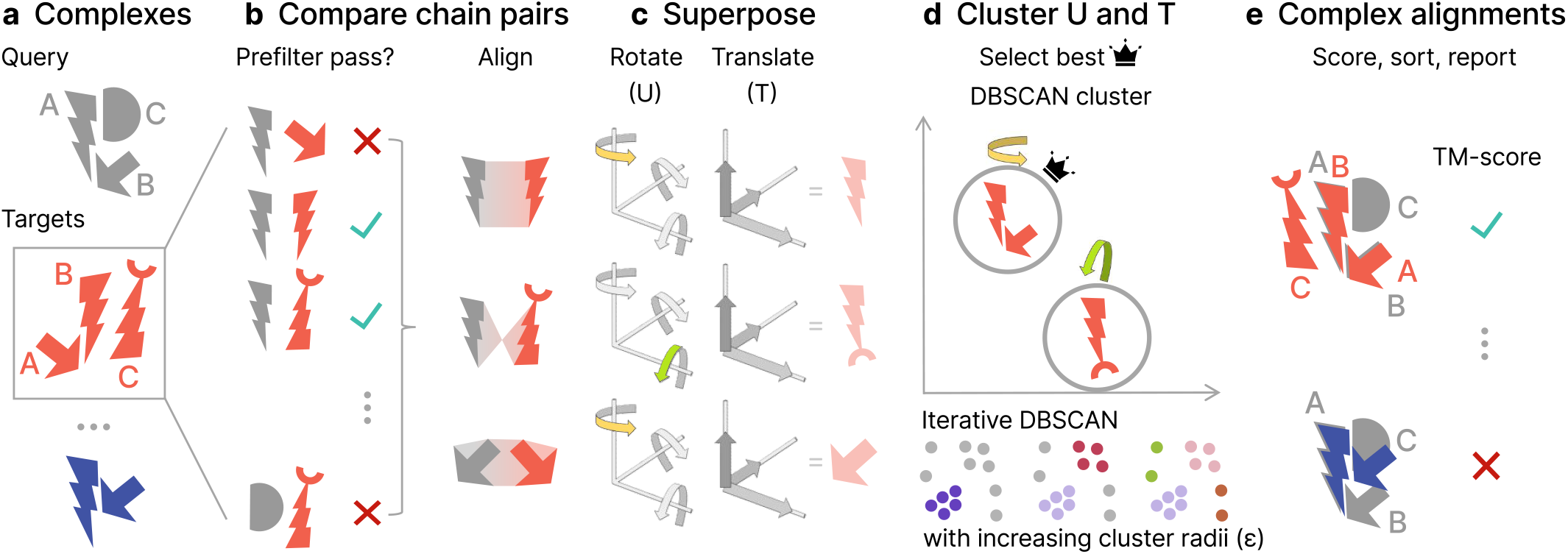
Foldseek-Multimer Schematic. **a**, Foldseek-Multimer allows fast querying of input complex(es) against a large database, potentially containing millions of targets. **b**, All chains from the query (gray) are compared to those of each target (red). A prefilter allows to quickly reject non-matching chain pairs so the full alignment is only applied to promising complex pairs. **c**, Foldseek-Multimer represents each chain-to-chain alignment as a superposition, described by rotations and translations, required for superposing the target chain onto the query. In this simplified example, two chain-to-chain alignments (top, bottom) are a rotation along one axis (yellow and green highlights), while one (middle) is a rotation along a different axis. **d**, The complex-to-complex alignment is inferred from chain-to-chain alignments as the superpositions of chain-pairs in the complex alignment are similar (“Algorithm: Overview”). Foldseek-Multimer uses the DBSCAN algorithm iteratively, with increasing radii, to identify superposition clusters and the best-scoring valid cluster for computing the complex alignment (**Supp. Fig. 1**). **e**, Based on the best-scoring cluster, the complex TM-score is computed across all chain alignments between query and target.

The quality of Foldseek-Multimer’s alignments was compared to that of US-align on a benchmark of 931 pairs of protein complexes, known to be structurally similar, using either tool to align them. Foldseek-Multimer was run in two modes, differing in the algorithm used for chain-to-chain alignment: 3Di+AA (*Foldseek-MM*) or TM-align (13) (*Foldseek-MM-TM*). Both tools detected the vast majority (> 95%) of pairs as similar (US-align: 97.6%, *Foldseek-MM-TM*: 97.4% and *Foldseek-MM*: 95.8%), aligning them with a TM-score ≥ 0.65, which is a cutoff found to be optimal for detecting structural similarity among complexes (6). Using either mode, Foldseek-Multimer computed highly correlated TM-scores to those of US-align (**Fig. 2a, Supp. Fig. 2**) and produced the same chain pairing in >99% of the cases (Data Availability).

**Fig. 2.**
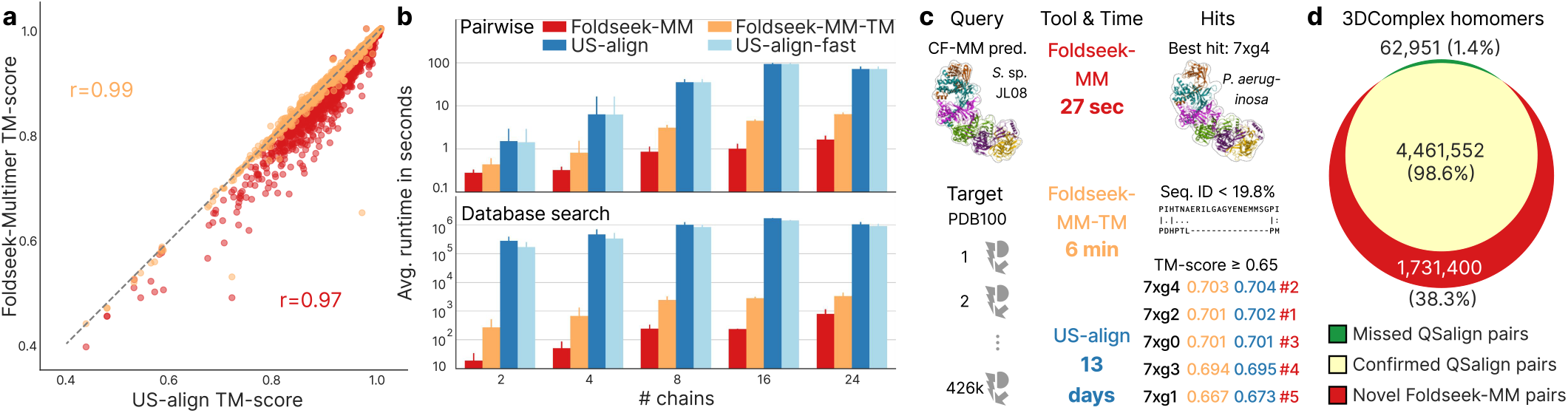
Performance of Foldseek-Multimer. **a**, Query-length normalized TM-scores (target-normalized: Supp. Fig. 2) computed for 931 pairs of structurally similar complexes by US-align (x-axis) or Foldseek-Multimer (y-axis). Both measures correlate highly (Pearson’s *r*). **b**, Execution time based on the dataset used for panel a. Complexes were binned by their number of chains, selected bins are shown (all bins: Supp. Fig. 3). Components’ contribution to speed: in pairwise mode (top), alignment (*Foldseek-MM*) and superposition clustering (*Foldseek-MM* and *Foldseek-MM-TM*) make Foldseek-Multimer 10-100 times faster than US-align. In database search (bottom), complexes were queried against 3DComplexV7. Foldseek-Multimer is further accelerated by using its prefilter, making it 10^3^-10^4^ times faster. **c**, An AlphaFold-Multimer prediction of a part of a CRISPR-Cas ribonucleoprotein from an environmental sample (top-left) was queried by Foldseek-Multimer and US-align against PDB100. *Foldseek-MM-TM* identified the same hits as US-align, while being >3,000 times faster. These hits were top-ranked by *Foldseek-MM* (red) with TM-score > 0.5. Non-aligned components of 7xg4 (top-right) are set as transparent. **d**, Foldseek-Multimer was run on 57 billion pairs of complexes from 3DComplexV7. It discovered nearly all homomeric pairs previously identified as similar by QSalign, and found an additional 1.7M homomeric pairs (Supp. Fig. 5).

We measured the runtime of the tools, breaking down the contribution of Foldseek-Multimer’s components to its speed. First, given the task of computing 931 pairwise alignments, we observed a speedup of 1-2 orders of magnitude over US-align (**Fig. 2b top, Supp. Fig. 3**), reflecting the efficiency of the chain-to-chain alignment (*Foldseek-MM*) and superposition clustering (*Foldseek-MM* and *Foldseek-MM-TM*). The performance of *Foldseek-MM-TM* thus highlights the key contribution of Foldseek-Multimer’s innovative use of superpositions as an alternative to US-align’s global alignment. Next, the tools queried each of the 677 complexes in the benchmark (Online Methods) against the 3DComplexV7 database (7). Here, Foldseek-Multimer was 3 to 4 orders of magnitude faster than US-align (**Fig. 2b bottom, Supp. Fig. 3**) due to an additional speedup by its prefilter.

Recently, Altae-Tran et al. (14) discovered the first CRISPR-Cas type IV-A system with a specified interference mechanism in an environmental sample of *Sulfitobacter* sp. JL08. Intrigued by their finding, we predicted a part of its ribonucleoprotein complex structure using ColabFold-AlphaFold-Multimer (9, 15). The prediction was of acceptable quality (0.564 pTM) and we provided it as a query to Foldseek-Multimer and US-align in a search against the PDB100 (On-line Methods). Foldseek-MM and Foldseek-MM-TM demonstrated remarkable efficiency in comparing a query consisting of six chains and spanning 1,843 amino acids against the 426,347 entries of PDB100. These comparisons took only 27 seconds and 6 minutes, respectively, on a single core of a server (23 and 96 seconds on an 8-core MacBook Pro). By contrast, it took US-align 13 days.

Here, in addition to its fast core-algorithm (**Fig. 1**), Foldseek-Multimer gained further acceleration since PDB100 is a clustered database, allowing it to search against the 343,785 representatives, instead of all entries, and to expand the search only within promising clusters (Online Methods). *Foldseek-MM-TM* and US-align scored five entries above 0.65. These entries were the top ranks by *Foldseek-MM*, scoring above 0.5 but below 0.65 (**Fig. 2c**). All five hits were from a recently reported type IV-A system in *Pseudomonas aeruginosa* (16), which belongs to a different class (Gammaproteobacteria) than that of the query (Alphaproteobacteria). When examining the best match, 7xg4, we found that Foldseek-Multimer could identify similarity, despite low sequence similarity (11.1-19.8% sequence identity and 19-33.3% sequence similarity using the BLOSUM62 substitution matrix) between the six subunit pairs of *Sulfitobacter* sp. JL08 and those of 7xg4. This provides further support for the previous identification of the *Sulfitobacter* sp. JL08 system as type IV-A and highlights the potential of Foldseek-Multimer for investigating protein complex structures predicted in distant organisms from environmental samples (**Supp. Fig. 4**: prediction quality effect). Next, we examined Foldseek-Multimer in an all-vs.-all setting, using the 3DComplexV7 database (7) as it had been previously analyzed in this setting using QSalign (Online Methods). QSalign relies on the time-consuming Kpax (17) structural alignment method, which prohibits it from conducting an exhaustive structural search. Thus, it first identified ca. 58 million pairs, which shared sequence similarity and then applied Kpax only to them, detecting ca. 4.5 million pairs of similar homomers (Online Methods: “QSalign Pairs”).

Using 128 cores, *Foldseek-MM* then queried the clustered 3DComplexV7 (Online Methods) against itself, examining 57 billion pairs in 11h. Applying the same TM-score >= 0.65 cutoff as QSalign, *Foldseek-MM* identified 98.6% of the homomeric pairs previously identified by QSalign and found an additional 1.7 million similar homomeric pairs: “Foldseek-MM Pairs” (**Fig. 2d**). We used US-align for evaluating a randomly-selected sample of 10% of the Foldseek-MM pairs (Online Methods). US-align confirmed 98.2% of the sampled pairs and rejected 1.8% (TM-score < 0.65). We thus conclude that over 1.6 million of the homomeric pairs are new discoveries by Foldseek-Multimer, owed to its ability to detect similar complex structures below the twilight zone (**Supp. Fig. 5**).

In addition to developing a command-line tool, we extended the Foldseek webserver to support Foldseek-Multimer and visualize its search results using the NGL viewer library (18). The webserver overlays chain-to-chain assignments by using translucently colored protein surfaces. Users can choose between Foldseek-Multimer’s alignment modes, and apply taxonomic filters, restricting the search to specific clades. To accompany the webserver with predicted structures, we organized 297,570 multimer predictions from several community efforts (10, 11, 19–21) into a single database. This BFMD resource (Online Methods) is available both within the web-server and for local use.

In conclusion, we presented a novel strategy for complex-to-complex alignment, which quickly identifies compatible sets of chain-to-chain alignments by their superpositions. Demonstrated here on protein complexes, the Foldseek-Multimer strategy can be extended to other modalities, such a RNA and DNA complex structures, given a way to align their individual subunits. Taken together, the unprecedented sensitivity and speed offered by Foldseek-Multimer make it an essential tool for investigating complex structures in the AlphaFold era.

## Online Methods

### Algorithm: Overview

Foldseek-Multimer examines all possible chain-to-chain pairings between the compared complexes, using Foldseek (Fig. 1b). It then uses the fact that a structural alignment between two complexes, Q and T, implies a superposition: a set of rotations and translations, which minimize the sum of squared distances between their aligned residue pairs (22). For simplicity, assume Q and T to be two structurally similar dimers, consisting of the chains Q_A, Q_B and T_A, T_B, where Q_A is similar to T_A and Q_B is similar to T_B. The physical meaning of the complex-level structural similarity is that Q_A is positioned and oriented relative to Q_B within Q in the same way that T_A is positioned and oriented relative to T_B within T. Thus, the same superposition, i.e., the same set of rotations and translations, would minimize the distance between Q_A and T_A as well as the distance between Q_B and T_B. In other words, all individual chain-to-chain superpositions (e.g., the one between Q_A and T_A) are equal to one another and to the complex-to-complex superposition. Therefore, a set of chain-to-chain alignments is compatible and can define a complex-to-complex alignment, only if all chain-to-chain superpositions computed from that set are equal. Foldseek-Multimer therefore computes for each chain-to-chain alignment a vector, representing its superposition (Fig. 1c). Next, it uses DBSCAN (23) for clustering these vectors to identify compatible sets of chain-to-chain alignments, which share the same superposition and define valid complex alignments (Fig. 1d, Supp. Fig. 1). Once complex alignments are identified, Foldseek-Multimer computes their TM-score (24) and reports them (Fig. 1e).

### Algorithm: Input

Foldseek-Multimer allows for searching one or more query protein complex structures against a target complex structure, a database of complex structures or a database of clustered structures. Structures can be provided in PDB/mmCIF format or as a Foldseek-formatted database. Formatting structures is possible using the `createdb` command.

### Algorithm: Chain-to-chain alignments

By utilizing Foldseek, Foldseek-Multimer offers two main modes for chain-to-chain structure comparison. The default mode, 3Di+AA, encodes structures as sequences over a 20-state 3Di alphabet, as fully described by van Kempen et al. (2). Additionally, chain-to-chain alignments can be computed using TM-align (13), which is a global, albeit slower alignment method. During database search, a prefilter, which is based on the 3Di+AA mode, allows for a fast removal of most chain pairs, continuing to compute chain-to-chain alignments only on promising candidates.

### Algorithm: Chain-to-chain superposition vectors

Given a chain-to-chain alignment, Foldseek-Multimer computes the superposition of the target chain onto the query chain, using nine rotations (U) and three translations (T). In preparation for aligning complex structure Q and complex structure T, Foldseek-Multimer creates a matrix with 12 columns, whose rows are the superposition vectors, computed from all chain-to-chain alignments, belonging to Q and T. The mean and the standard deviation (SD) of each column are then used to compute the coefficient of variation (CV = SD/mean) of the column and exclude less-informative columns (CV < 0.1, Supp. Table 1: the effect of this parameter on Foldseek-Multimer’s performance). If the mean value of the column is < 1, the SD value is used instead of the CV for the exclusion criterion. Finally, the retained columns undergo normalization since they can have different scales. To that end, Foldseek-Multimer subtracts from each column its mean and divides it by its SD. We denote the resulting reduced and normalized matrix as *supQT*.

### Algorithm: Chain-to-chain clustering

DBSCAN is used iteratively for clustering the rows of *supQT* as it doesn’t require knowing the number of clusters *a priori*. The stages of this procedure are described below and demonstrated on a small example in Supp. Fig. 1.

#### Initialization

The Euclidean distances between all row pairs in *supQT* areis computed and the minimum (minDist) initializes the parameter epsilon. The biggest cluster(s) encountered during the procedure are recorded in a candidate list alongside their size (maxClusterSize), which is initialized to 0.

#### The DBSCAN iteration

For each *supQT* row, all rows within a radius of epsilon from it, are defined as its *neighbors*. Then, all rows, which have at least one more neighbor (at least two neighbors, including itself) are considered as “core-points” and the rest as “non-core-points”. Next, a core-point is selected at random to start the first cluster. All its core-point neighbors are added to the first cluster. Each added core-point neighbor also adds its core-point neighbors and so on, until no more core-points can be added to the first cluster. Then all non-core-points, which are neighbors of members of the first cluster, are added to it as well (without adding their neighbors). The second cluster is constructed similarly, operating on the remaining unclustered points.

#### Cluster validity and rescuing by Nearest Neighbors

During the DBSCAN iteration, after each cluster is computed, Foldseek-Multimer evaluates its validity. If a cluster includes the same chain in multiple chain-to-chain alignments, Foldseek-Multimer attempts to rescue it by selecting a compatible subgroup of points (i.e., chain-to-chain alignments) from that cluster. To that end, points are selected for the subgroup in the order of their distance to the core-point, which was used to initiate the cluster. Selection for the subgroup is stopped once the process encounters a point that includes a chain, which was already added by a previous point.

At each DBSCAN iteration, valid clusters which are at least as big as maxClusterSize are added to the candidate list. The value of maxClusterSize is updated each time a bigger cluster is encountered and all previously-added clusters are removed from the list due to being smaller.

#### Iterativity

Next, the value of the radius epsilon is increased by a delta of 0.1 (Supp. Table 1: the effect of this parameter on Foldseek-Multimer’s performance) and a new DBSCAN iteration starts, potentially forming new clusters. If all new clusters are smaller than maxClusterSize the procedure stops. Otherwise, the candidate list of and maxClusterSize will be updated with the iteration’s clusters and epsilon will increase again, up to a maximal value of the distance between the two furthest points (maxDist).

#### Early stop condition

Let *C*_*Q*_ and *C*_*T*_ be the number of chains in Q and T, respectively. Without loss of generality, assume *C*_*Q*_ < *C*_*T*_. If maxClusterSize is equal *C*_*Q*_, then no bigger valid cluster exists. Since there is a total of C_Q_xC_T_ chain-to-chain alignments, the number of clusters in the candidate list cannot exceed *C*_*T*_ once maxClusterSize is equal to *C*_*Q*_. Foldseek-Multimer checks these two conditions and avoids unnecessary DBSCAN iterations if they are met.

#### Discovered clusters

At the end of the iterative DBSCAN procedure, the biggest valid clusters are returned. Each of them is equivalent to a set of compatible chain-to-chain alignments with a similar superposition that together define a complex alignment between Q and T.

### Algorithm: TM-score computation

TM-scores are computed for the complex alignment derived from each of the valid clusters found for a Q-T complex pair as follows. First, the chains of complex Q are concatenated to each other in some order. Given the concatenation order of the chains in Q, Foldseek-Multimer concatenates the chains of complex T, in the order of their pairwise matches to the chains of Q, as defined by the cluster. Then, the TM-score between the concatenated Q and concatenated T is computed the same way Foldseek computes it for single-chain pairwise alignments, using the C*α* coordinate vectors of both chains (concatenated chains in this case). Using this computation, all complex alignments a given query complex Q has with a specific target T and with all other target complexes can be ranked and reported by their TM-score.

### Algorithm: Utilizing clustered databases

In order to further accelerate Foldseek-Multimer, we aimed to reduce the redundancy in the target database, an approach, which is also adopted by TM-search (25). To that end, we introduced a new capability to Foldseek, which allows it to efficiently search through clustered databases in MMseqs2 or Foldseek format (e.g., PDB100, see section). If the input has M cluster representatives and N cluster members (M < N), Foldseek will first search (prefilter + alignment) against the M representatives, finding candidates below a specific E-value threshold (the default value of 10 was used in this study). Extending to promising clusters only, the alignment step will then be carried out on all cluster members of the candidates. Foldseek-Multimer will use the alignment results of all extended clusters for computing superposition matrices and the following procedure steps, as described above.

### The 3DComplex database and QSalign comparisons

For the analyses presented in Fig. 2a, 2b and 2d, we downloaded the 3DComplex database version 7 (3DComplexV7 DB; Data Availability). Briefly, this database holds 238,965 structures, consisting of 557,146 chains and was created from the “Biological Units/Assemblies” downloaded from the PDB using the method described by Levy et al. (7). Prior to this study, QSalign (6) had been applied to 3DComplexV7 DB and yielded a list of 57,953,513 compared structural pairs.

#### Similar pairs benchmark

#### Dataset

Starting with the list of 57,953,513 QSalign-compared pairs, we selected entries with varying numbers of subunits (from 2 to 24). For each size, the criteria for selection were that the TM-score computed by Kpax (17) was greater than 0.8, and that pairs of homomers had less than 80% sequence identity. If more than 100 pairs matched the criteria, only the first 100 were selected, resulting in a total of 931 complex pairs included in the benchmark.

#### Runtime evaluation

Performance was measured on a server with a 1x AMD EPYC 7702P 64-core CPU and 1 TB RAM, using a single core. The queries for the time measurements in Fig. 2b and Supp. Fig. 3 were the 677 unique complexes associated with the 931 pairs. Due to its high computational demand, the runtime of US-align on these 677 complexes against 3DcomplexV7 was extrapolated from running against 1,000 randomly-sampled 3DcomplexV7 entries. Reporting the average over the number of cases Nc = 142,109,124,18,101,7,42,8,41,44,17,5,5,14 for each number of chains c = 2,3,4,5,6,7,8,9,10,12,14,16,18,24: 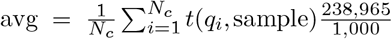. Foldseek-Multimer was run against the full database, without extrapolation.

### Environmental CRISPR-Cas

#### The PDB100 database

A version of the PDB, termed PDB100 was used to search for structural homologs of an environmental CRISPR-Cas as well as to measure the runtimes of Foldseek-Multimer and US-align. PDB100 was first introduced by van Kempen et al. (2), but further developed in this study, as described here. First, PDB, containing the asymmetric unit of 207,937 entries, consisting of 1,047,615 chains, was downloaded in November 2023 (Data Availability). Of these, 11,901 entries were associated with more than one structural model (e.g., the NMR experiment 2KOX). In total, 426,347 structural models were associated with the PDB entries. Next, all chains were clustered using Foldseek (parameters: `-c 0.95 --min-seq-id 1.0`), resulting in 343,785 redundancy-reduced representatives. In contrast to van Kempen et al., PDB100 is now a cluster database, which holds the representatives alongside information to associate them to their cluster chains and structural models. PDB100 is updated regularly and is available through the Foldseek webserver and can be downloaded using the *databases* command.

#### Complex structure prediction

Four *Sulfitobacter* sp. JL08 protein sequences, identified as CRISPR-Cas type IV-A components by Altae-Tran et al. (14): Csf1, Csf2, Csf3 and Cas6 were obtained from the plasmid map “pHS1068 NZ_CP025815 DinG HNH proteins (E. coli codon optimized) CRISPR array in pACYCDuet-1 with Lac promoters.gb”, released by the authors. Following the reported stoichiometry of the CRISPR-Cas type IV-A core complex (26), we constructed an input file for ColabFold-AlphaFold-Multimer (15) with eight chains: 1xCsf1 + 5xCsf2 + 1xCsf3 + 1xCas6. When examining the structure, we noticed that AlphaFold-Multimer did not predict an interaction between Csf1 and Cas6 and the rest of the complex, so we omitted them and re-predicted the structure: 5xCsf2 + 1xCsf3. Comparing the four sequences of *Sulfitobacter* sp. JL08 to protein nr (27) was performed using the blastp webserver (Feb. 2024).

#### Runtime evaluation

Performance was measured on a server with a 1x AMD EPYC 7702P 64-core CPU and 1 TB RAM, using a single core. Due to its high computational demand, the total runtime was extrapolated when measuring US-align on the *Sulfitobacter* sp. JL08 structure against the PDB100, using five samples: 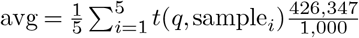. Foldseek-Multimer was run against the full database, without extrapolation. For the MacBook runtime measurements, we used a 13-inch Macbook Pro (M1; 2020; Model A2338) with 16GB RAM.

### Comparison to QSalign on 3DComplex V7

#### QSalign Pairs

Starting with the list of 57,953,513 QSalign-compared pairs (see section), high-scoring homomeric pairs (max TM-score >= 0.65) were selected, excluding pairs with a PISA structure. This resulted in 4,524,503 structurally similar unique homomeric pairs, which we denoted “QSalign Pairs”.

#### A clustered 3DComplex V7

The 557,146 chains of 3DComplex V7 (see section) were clustered using Foldseek (parameters: `-c 0.99 --min-seq-id 0.9 -e 0.00001`), resulting in 142,957 redundancy-reduced representatives. This procedure took 18 seconds, using 64 threads.

### The BFMD resource

In an effort to generate a large multimer database, we gathered 297,570 multimer predictions, consisting of 597,640 chains from several community efforts. These were turned into a clustered Foldseek database using the parameters: `-c 0.95 --min-seq-id 1.0 -e 0.00001`, resulting in 51,757 redundancy-reduced representatives. All predictions’ accessions are prefixed by the resource name. Multimers extracted from the `ModelArchive` (19), all-vs-all prediction of a set of human genome maintenance proteins `Predictomes` (20), `LevyLab` atlas of predicted homomers (10), protein-protein prediction from the Human Reference Interactome (28) and the Human Protein Complex Map (29) `HuIntAF2` (11) and `ProtVar`, predicted multimers as part of an effort to understand missense variance (21). The BFMD is available through the Foldseek webserver and is downloadable as a standalone database using the `databases` module.

#### Foldseek-MM all-vs.-all search of 3DComplex V7

During this search all temporary files were kept in memory and 128 cores were used (2 x AMD EPYC 7742). The entire search finished in 10 hours and 23 minutes. Most of the time was spent in the module for matching chains, which took 7 hours and 32 minutes.

#### Evaluation of “Foldseek-MM Pairs”

About 1.7 million pairs of homomeric complexes were detected only by *Foldseek-MM* as similar. Since running US-align over all pairs is prohibitively slow, we randomly selected 160,252 pairs (ca. 10% of all pairs) and computed their alignment using US-align. For 2,844 of these (1.8%), US-align reported a TM-score < 0.65, which we use as an estimate for the false-positive rate among the full set of novel “Foldseek-MM Pairs”. 157,391 pairs (98.2%) were confirmed as matches by US-align and the rest (17 pairs, <0.0001%) were aligned as monomers.

### Tool commands and arguments

*Foldseek-MM* commit c27a629 (default, using 3Di+AA):

~~~
foldseek easy - complexsearch query. pdb
target. pdb / target DB result tmp -- threads 1
~~~

*Foldseek-MM-TM* commit c27a629 (using tmalign):

~~~
foldseek easy - complexsearch query. pdb
target. pdb /target DB result tmp -- threads 1
-- alignment - type 1
~~~

Additionally, the flag ‘`--exhaustive-search` 1’ was used for the benchmark of similar pairs and the flag ‘`--cluster-search` 1’ when using a clustered db. For database searches we pre-indexed the database using ‘createindex targetDB’ and kept it in memory. We set `--db-load-mode` to 2 in easy-complexsearch, to indicate that the pre-indexed database is already in memory. During database search, Foldseek-Multimer can include or reject monomeric targets in the reference database using the `--monomer-include-mode` parameter. For this study, we set the parameter to reject all monomer matches.

*US-align* version 20220924:

~~~
US - align query. pdb target. pdb - mm 1 - ter 0 - mol prot
~~~

Additionally, the flag ‘`-fast`’ was set for during runtime assessments in Fig. 2b). For speed measurements we kept the PDB/mmCIF files in memory to avoid IO related bottlenecks.

## Data availability

The benchmarking and 3DComplexV7 data is available at https://doi.org/10.5281/zenodo.13121434. PDB: files.wwpdb.org/pub/pdb/data/structures/all

## Code availability

Foldseek-Multimer and its webserver are GPLv3-licensed free open-source software. The source code and binaries for Foldseek-Multimer can be downloaded at github.com/steineggerlab/foldseek. The analysis scripts are available at github.com/steineggerlab/foldseek-multimer-analysis. The webserver is available at search.foldseek.com and its source-code at github.com/soedinglab/mmseqs2-app.

## Acknowledgments

We thank Hyunbin Kim for helping with designing Figure 1, Dr. Yong Hwan Kim for suggesting the name Foldseek-Multimer, and Dr. David Burstein for his feedback on the CRISPR-Cas example. We thank the EMBL-EBI for releasing the “EMBL-EBI Icon fonts for the life sciences” that are used in the webserver. M.S. acknowledges support by the National Research Foundation of Korea grants (2020M3-A9G7-103933, 2021-R1C1-C102065, 2021-M3A9-I4021220 and RS-2024-00396026), Samsung DS research fund, Creative-Pioneering Researchers Program and AI-Bio Research Grant through Seoul National University. M.M. acknowledges support from the National Research Foundation of Korea (grant RS-2023-00250470). E.L. acknowledges support from the European Research Council (ERC) under the European Union’s Horizon 2020 research and innovation program (grant agreement No. 819318).

## Conflict of interest

M.S. acknowledges outside interest in Stylus Medicine.

## Author contributions

W.K., J.S. and M.S. designed the Foldseek-Multimer algorithm. W.K., M.M. and M.S. developed the software. M.M. and C.G. developed the Foldseek-Multimer webserver. E.L. and H.S. designed the comparison to QSalign, provided 3DComplexV7 and applied QSalign to it. E.L.K. developed the CRISPR-Cas example. W.K., E.L.K., and M.S. designed the figures, performed the benchmarks and wrote the manuscript, with contributions from all authors.

**Supplementary Table 1.** Effect of parameters on Foldseek-Multimer sensitivity and runtime. Foldseek-Multimer uses two hard-coded parameter values: epsilon-delta = 0.1 and CV = 0.1 (Online Methods, “Algorithm”). Here, we tested the impact of alternative values for these parameters on Foldseek-Multimer’s sensitivity and runtime. Sensitivity was measured by fitting a linear regression model between US-align’s TM-scores (the independent variable *X*) and Foldseek-Multimer’s TM-scores (the dependent variable *Y*) computed for the dataset of 931 structurally-similar complex pairs of Fig. 2a. We then examined the estimates for the coefficients of the model *Y* = *b*0 + *b*1 × *X* as well as the correlation between *X* and *Y* (expressed as Pearson’s *r*) for each alternative parameter setting compared to the default (first row). Lower (but not higher) values of the DBSCAN parameter epsilon-delta have the potential of improving sensitivity, however we did not observe any improvement over the default in either setting, with similar runtimes (averaged over 5 runs). We tested two more conservative CV cutoffs for the superposition matrix column exclusion, which resulted in worse correlations between Foldseek-Multimer and US-align compared to the default. This suggests a lower CV cutoff introduces more noise and is therefore less preferable. We also tested a higher value of CV, which resulted in the same correlation as the default. Taken together, these results support the choice of the hard-coded parameter values.

| Epsilon-delta | CV | Pearson's $r$ | $\hat{b}_0$ | $\hat{b}_1$ | Avg. runtime (s) |
| --- | --- | --- | --- | --- | --- |
| 0.1 | 0.1 | 0.97 | -0.124 | 1.107 | 605 |
| 0.05 | 0.1 | 0.97 | -0.124 | 1.107 | 601 |
| 0.01 | 0.1 | 0.97 | -0.124 | 1.107 | 603 |
| 0.1 | 0.05 | 0.94 | -0.119 | 1.101 | 600 |
| 0.1 | 0.01 | 0.85 | -0.129 | 1.106 | 591 |
| 0.1 | 0.2 | 0.97 | -0.124 | 1.107 | 600 |

**Supplementary Figure 1.**
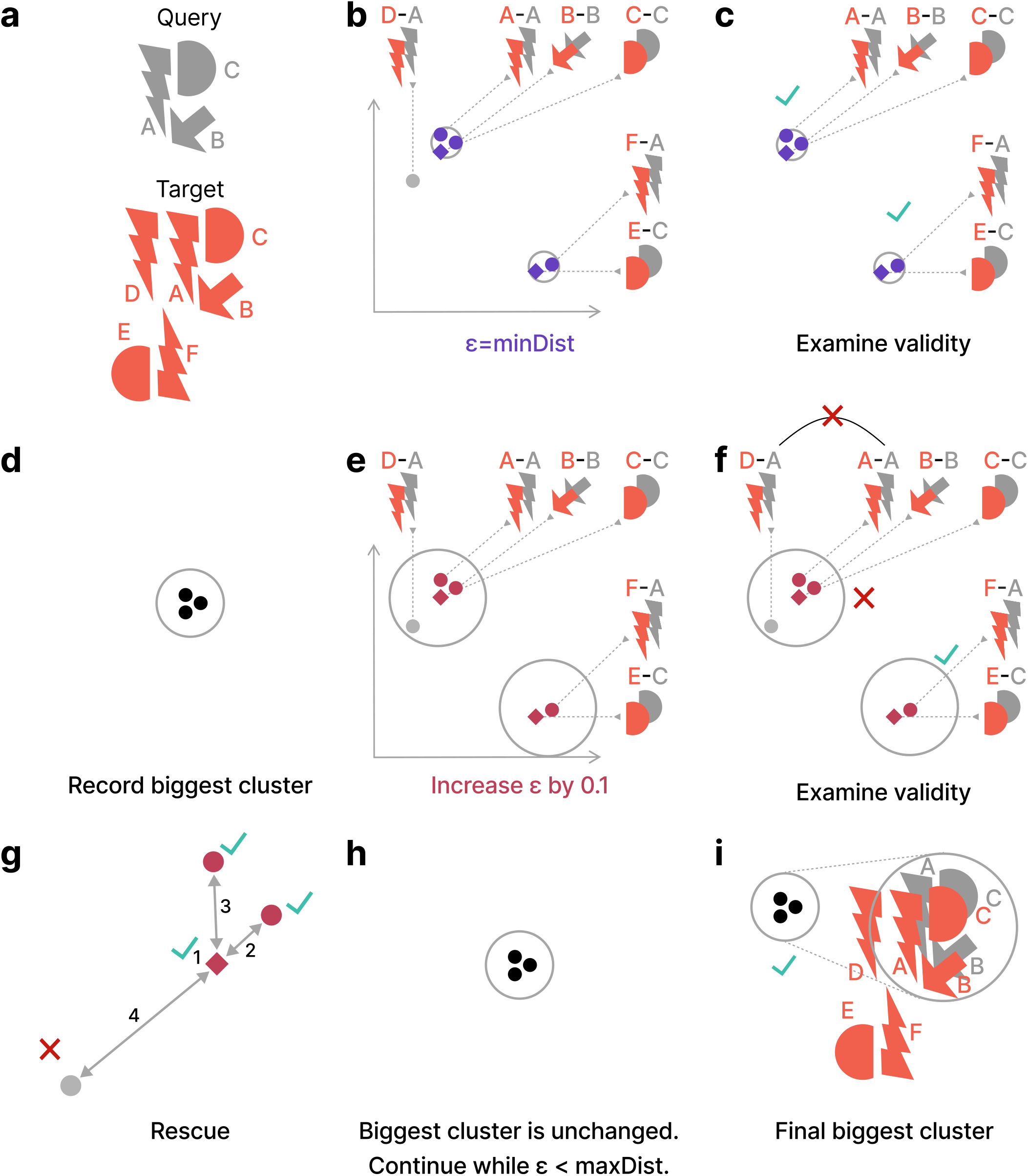
Iterative DBSCAN procedure for clustering chain-to-chain alignments. The procedure steps are demonstrated schematically on a small example and fully described in the Online Methods section “Algorithm: Chain-to-chain clustering”. **a**, A query complex can match a target complex in two ways. The first, in which chains Q(A,B,C) are matched to T(A,B,C) is the best match with a certain superposition. The second, in which Q(A,C) are matched to T(F,E) is a sub-optimal match with a different superposition. In addition, query chain A can match target chain D with a third superposition, which is somewhat close to the best superposition (but not identical). **b**, after the chain-to-chain superpositions (points) are computed they are clustered using DBSCAN with *ϵ* = *minDist*, the minimal distance between any two points. Five points have enough neighbors to be considered as core-points (purple) and two clusters are discovered. The core-points from which the clusters started are marked as diamonds. **c**, Both clusters are valid as they contain at least two chains and no chain is repeated. **d**, The biggest (most chains) cluster is recorded. **e**, *ϵ* is increased by 0.1 and a new DBSCAN clustering begins with core-points (burgundy) and discovers two clusters. **f**, One of the clusters is not valid since query chain A is matched twice to chains A and D on the target. **g**, In a rescue procedure, the points of the cluster are examined by their distance (numbered arrows) to the starting core-point (diamond), collecting a valid subset of points and stopping when a violating point is reached. **h**, The biggest valid cluster is unchanged in this example. The procedure continues until *ϵ* is greater than the distance between the two furthest points (maxDist). **i**, The final biggest cluster(s) discovered over all iterations is used for defining the complex-to-complex alignment and computing its TM-score.

**Supplementary Figure 2.**
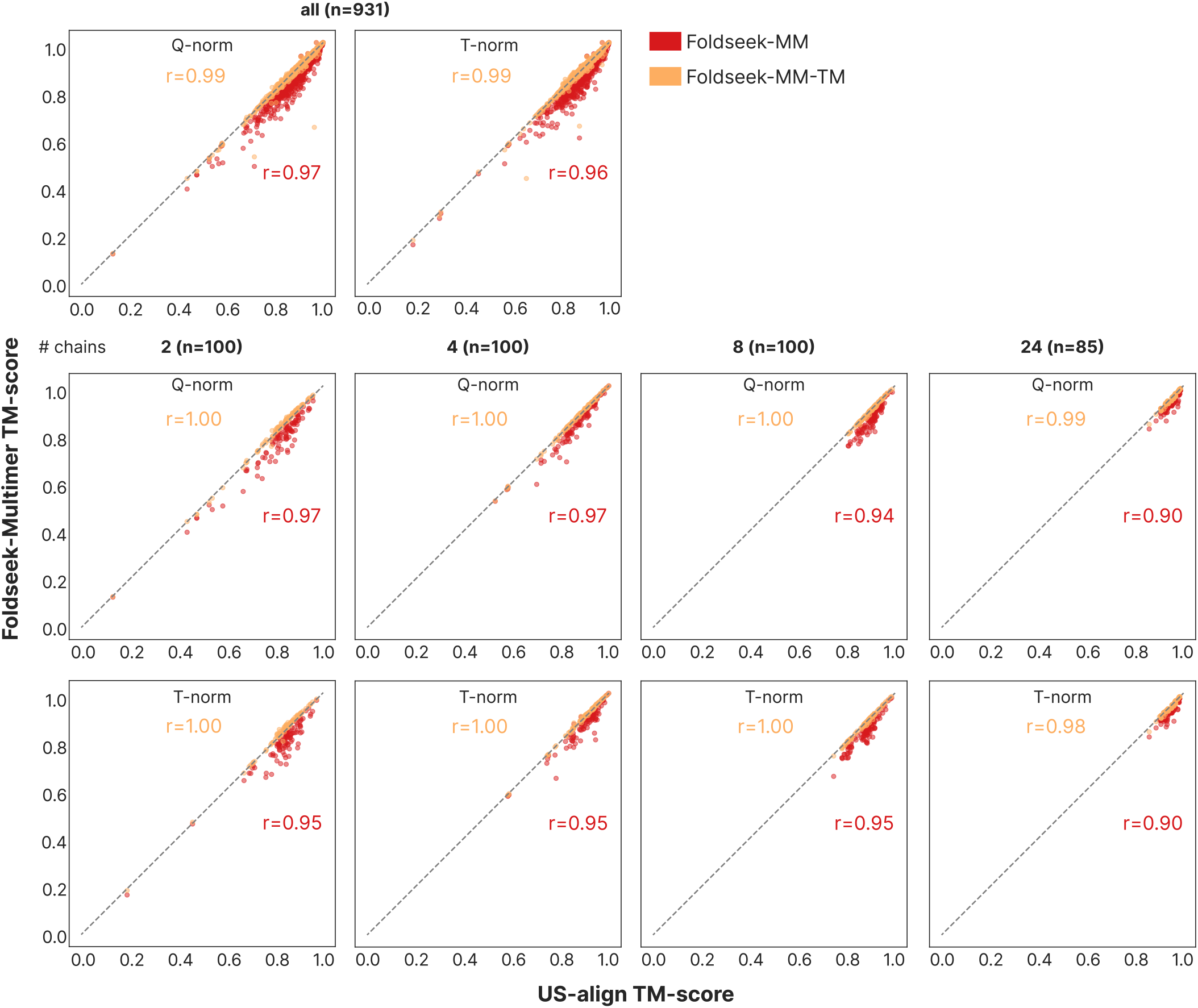
TM-score correlation by normalization method and number of chains. Fig. 2a presents the correlation between query-length normalized (Q-norm) TM-scores computed by either Foldseek-Multimer or US-align for 931 pairs of structurally similar complexes. This computation was repeated using target-length normalization (T-norm), resulting in similarly very high correlation between the tools (top panel, right). In addition, the correlation was computed separately for complexes of the same number of chains using either query- or target-length normalization if there were at least 30 complexes of that size (i.e., separately for complexes with 2, 3, 4, 5, 6, 8, 10, 12, 14, and 24 chains). For all computed cases, the correlation was found to be high (Pearson’s *r* : 0.90-1.0). The middle and bottom panels depict these correlations for a subset of the separately-computed cases. The sample size n is indicated in parentheses.

**Supplementary Figure 3.**
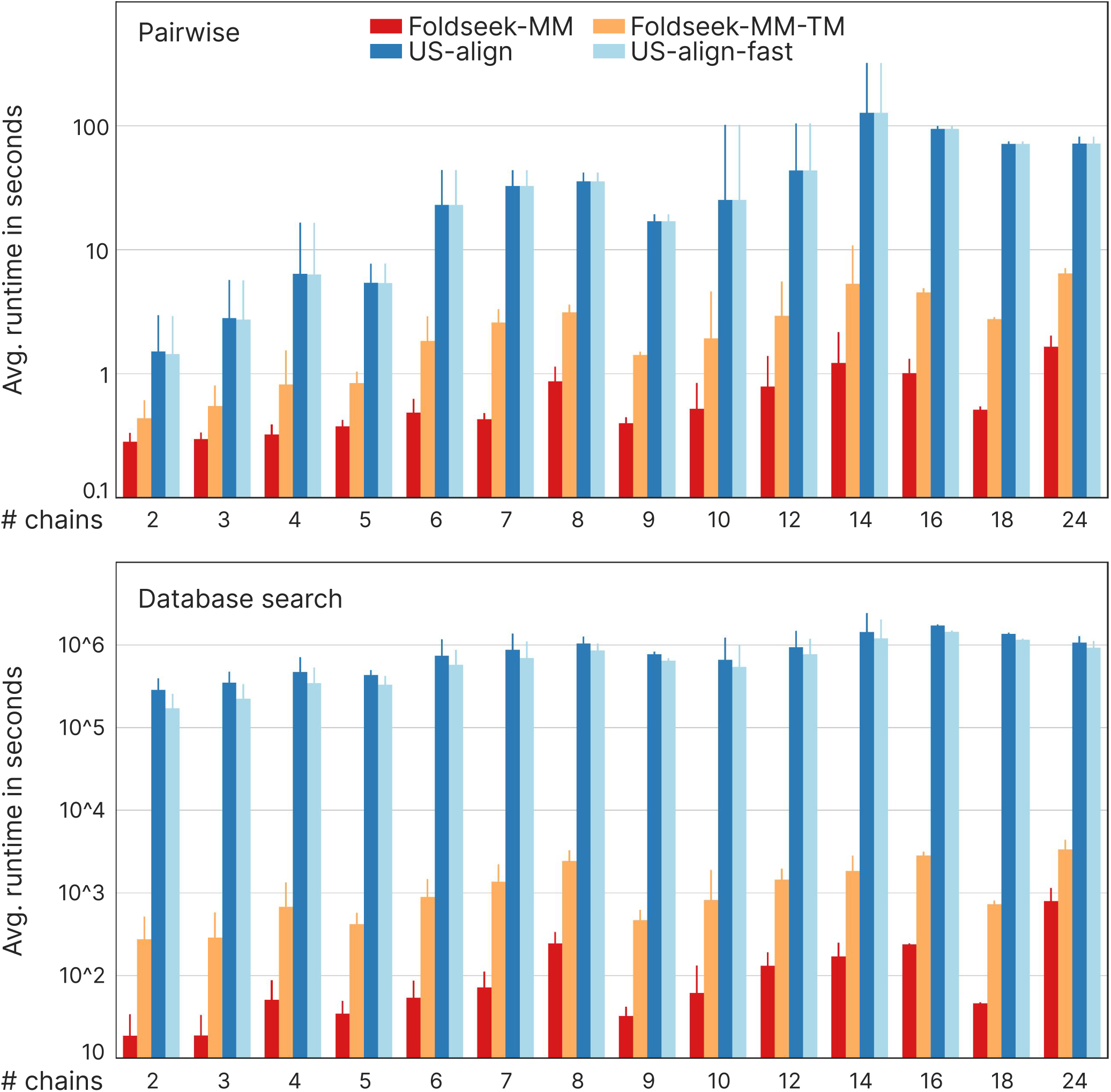
Speed comparison of Foldseek-Multimer to US-align. Execution time based on the dataset used for Fig. 2a-b. Complexes were binned by their number of chains. Speed comparison of pairwise alignment (top): bar height depicts the average and standard error computed for each bin over the following number of cases: 100, 100, 100, 43, 100, 22, 100, 18, 82, 100, 51, 18, 12, and 85. Foldseek-Multimer is ×10 − 100 faster than US-align due to efficient chain-to-chain alignment and superposition clustering. Speed comparison of Database search (bottom), complexes were queried against 3DComplexV7: bar height depicts the average and standard error computed for each bin over the following number of cases: 142, 109, 124, 18, 101, 7, 42, 8, 41, 44, 17, 5, 5, and 14. Foldseek-Multimer is further accelerated by its prefilter, making it ×10^3^-10^4^ faster.

**Supplementary Figure 4.**
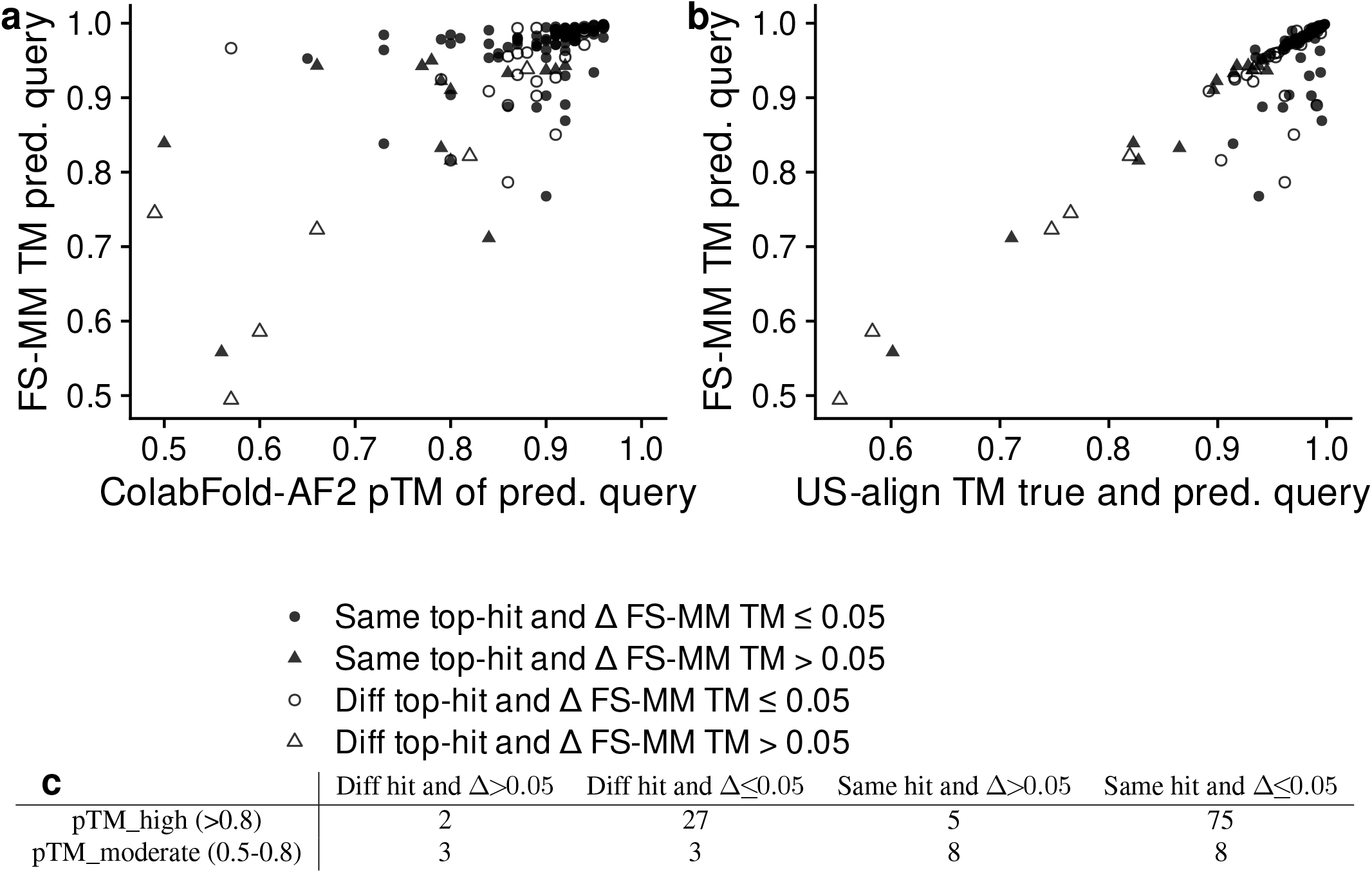
Effect of complex structure prediction quality on Foldseek-Multimer. The results presented in Fig. 2a are based on PDB structures, which have been determined using gold-standard techniques and can be considered as “true”. Here, we sought to measure the effect of using predicted query structures, as would be the case in metagenomic studies, like the analysis presented in Fig. 2c. To that end we used ColabFold-AlphaFold2 v1.5.5 to predict 50 (5 models ×10 seeds) structures for each of the 132 unique dimers, which were shorter than 1,000 residues, from the Fig. 2a dataset. We set the prediction parameters to produce structures of various qualities (`--use-dropout --num-seeds 10 --num-recycle 6`). For each of the dimers, we took the lowest ranked prediction. The quality of the predicted queries was measured in two ways: by the pTM score, estimated by ColabFold-AlphaFold2 and by the US-align TM-score between the true and predicted structures. While the latter is more accurate, this information will not be available in a metagenomic study, where the true structure is unknown. We then used Foldseek-Multimer to search each predicted query complex against the PDB100 database, recorded its top hit and score, and compared it to the top hit and score recorded for the true structure. **a and b**, higher quality predictions tend to result in better FS-MM TM-scores. At the same time, despite the variance in prediction quality, all but three (97.7%) predicted structures had a FS-MM top hit with a TM-score > 0.65. Of these, most (85.6%) had the a very small FS-MM TM-score difference (≤ 0.05) when querying with the predicted and the true structure (circles). **c**, Predicted structures that were less similar to the true one (pTM_moderate) were more likely to have a worse score (triangles, Δ = FS-MM score true query - FS-MM score predicted query) compared to well predicted structures (pTM_high). A chi-square test of independence was performed to examine the relation between the quality of the predicted structure and the difference in score. This association is statistically significant (Chi-square statistic = 29.3, N = 131, DF = 1, P-value < 0.00001).

**Supplementary Figure 5.**
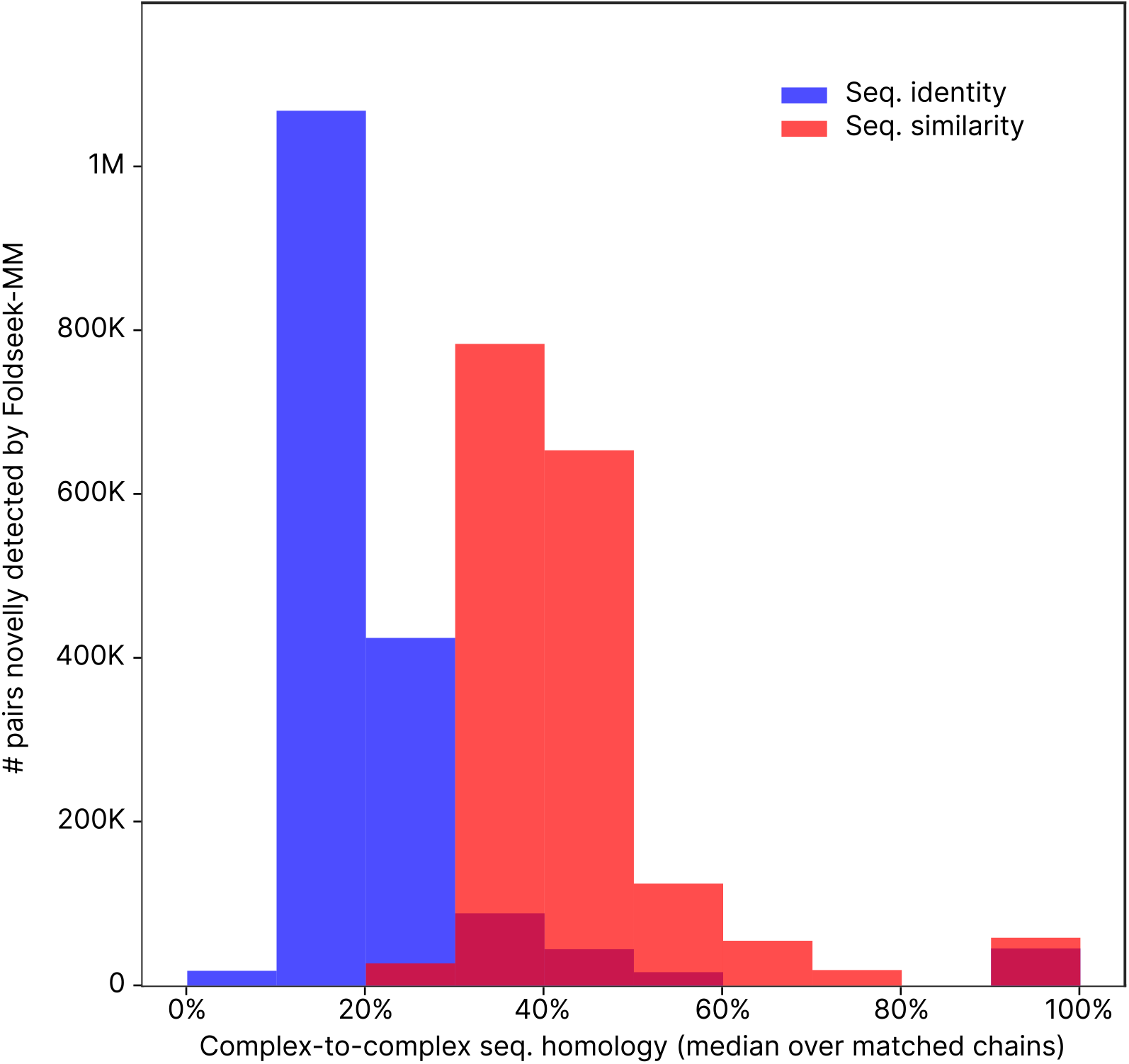
Comparison of 3DComplexV7 complex pairs novelly discovered by Foldseek-MM. When applied to the 3DComplexV7 database, Foldseek-MM novelly reported 1,731,400 pairs of homomeric complexes as structurally similar (**Fig. 2d**). For these pairs we measured the median sequence identity (blue) or similarity (red, computed using the BLOSUM62 substitution matrix) over all matched chains of that pair. Most complex pairs (85%) were highly diverged, having less than 50% sequence similarity. Furthermore, these results demonstrate Foldseek-Multimer’s ability to detect structural similarity well below the twilight zone as 87% of the pairs had less than 30% sequence identity between the complexes.

## Notes

https://github.com/steineggerlab/foldseek

https://search.foldseek.com/multimer

### Summary of Updates

We improved the speed of Foldseek-Multimer. Fixed numbers in the text. Updated figures.

